# Regional missense constraint improves variant deleteriousness prediction

**DOI:** 10.1101/148353

**Authors:** Kaitlin E. Samocha, Jack A. Kosmicki, Konrad J. Karczewski, Anne H. O’Donnell-Luria, Emma Pierce-Hoffman, Daniel G. MacArthur, Benjamin M. Neale, Mark J. Daly

## Abstract

Given increasing numbers of patients who are undergoing exome or genome sequencing, it is critical to establish tools and methods to interpret the impact of genetic variation. While the ability to predict deleteriousness for any given variant is limited, missense variants remain a particularly challenging class of variation to interpret, since they can have drastically different effects depending on both the precise location and specific amino acid substitution of the variant. In order to better evaluate missense variation, we leveraged the exome sequencing data of 60,706 individuals from the Exome Aggregation Consortium (ExAC) dataset to identify sub-genic regions that are depleted of missense variation. We further used this depletion as part of a novel missense deleteriousness metric named MPC. We applied MPC to *de novo* missense variants and identified a category of *de novo* missense variants with the same impact on neurodevelopmental disorders as truncating mutations in intolerant genes, supporting the value of incorporating regional missense constraint in variant interpretation.

## Introduction

With the widespread accessibility of genome sequencing, interpreting genetic variation has become a central challenge in medical genetics. Particularly given that the vast majority of variants in any given genome are benign, the ability to pinpoint true disease associated variation within a vast background of neutral variation is key to both discovery and diagnostics. This has specifically arisen in the study of *de novo* (newly arising) variants. Over the last few years, many projects sequenced thousands of parent-child trios—primarily focused on neurodevelopmental disorders, such as intellectual disability^1^^-^^3^, developmental delay^4,5^, epileptic encephalopathy^6^, and autism spectrum disorders^7^^-^^12^—to evaluate the role of *de novo* variation in the hope of identifying genes and pathways relevant to disease etiology. These studies established and characterized the important, but limited, role of *de novo* variation in the genetic architecture of these diseases. The largest excesses were found for *de novo* protein-truncating variants (PTVs) (roughly 2-fold enriched over expectation; **Table S1**), which because of their infrequency and obvious functional consequence have become the main focus for follow up research.

The availability of large-scale exome sequencing datasets of reference individuals, such as the Exome Aggregation Consortium (ExAC; n = 60,706)^13^, enable refined searches for *de novo* PTVs most likely to contribute to disease. These databases provide the opportunity to better understand patterns and rates of variation within the human population and have permitted the identification of genes and regions within genes that are intolerant of nonsynonymous variation (constrained)^14^^-^^16^ and therefore more likely to be associated to disease. As an example, Kosmicki and colleagues found that the overall 2.6-fold excess for *de novo* PTVs in patients with neurodevelopmental disorders (when compared to controls) is explained almost entirely by a small subset of variants, specifically those *de novo* PTVs, absent from individuals in ExAC, which disrupted genes which were recognizably and strongly intolerant of lossof-function variation (6.7-fold enriched; no significant signal in the remaining *de novo* PTVs)^17^.

Beyond PTVs, cases with neurodevelopmental disorders also have a significant, but more modest (1.3-fold; **Table S1**), enrichment of *de novo* missense variants compared to expectation, indicating that a subset of these variants are disease-related. Interpreting missense variation, however, presents unique challenges. While some amino acid substitutions may lead to effects on the protein just as deleterious as PTVs (or in the case of some gain-of-function substitutions, even more so), many substitutions can be neutral. Additionally, there are position-dependent effects of missense variants, meaning that neutral substitutions can occur next to disease-relevant substitutions. This stands in contrast to PTVs where, for the most part, truncating variants at adjacent amino acids have the same effect (i.e., it makes little difference if a transcript is truncated at 20% or 50% of full-length if both are eliminated by nonsense-mediated decay). These two elements—the location and specific amino acid substitution—make interpreting missense variation considerably more challenging than interpreting PTVs.

Previous tools used to predict the deleteriousness of missense variation have relied primarily on sequence conservation across species (e.g. SIFT^18^), structural features of the protein (e.g. PolyPhen-2^19^), or combinations of these metrics (e.g. CADD^20^, MutationTaster^21^). Few metrics have yet to take advantage of the knowledge of constrained genes and regions, where natural selection most aggressively removes variation within the human population. One metric to do so is M-CAP^22^, which includes a metric of genic intolerance to nonsynonymous variation (RVIS^14^) as one of the many features in its classifier. Given that the deleteriousness of missense variation is tied to the domain disrupted by the variant, we hypothesize that incorporating local depletion of missense variation will increase our ability to differentiate pathogenic from benign variation.

To pursue this idea, we developed a method to identify regions within genes that are specifically depleted of missense variation and found that ~15% of genes show evidence of regional variability in missense intolerance (constraint). That is, genes with two or more sub-regions with significantly different tolerance for missense variation. We then combined this information with variant-level metrics to best predict the impact of any observed missense variants. There are many tools to predict the deleteriousness of missense variants^19^^-^^22^ and to evaluate specific amino acid substitutions^23,24^. To this end, we created a score (missense badness) that measures the increased deleteriousness of amino acid substitutions when they occur in missense-constrained regions. We combined information from orthogonal deleteriousness metrics into one score, called MPC (for **M**issense badness, **PolyPhen-2**, and **C**onstraint). When we evaluate the MPC scores of *de novo* missense variants, we found that 17.6% of missense variants from neurodevelopmental disorder cases, but only 3.9% from controls, have MPC ≥ 2. Using MPC therefore allowed us to identify a subset of *de novo* missense variants with an effect size approaching that of *de novo* PTVs in intolerant genes, cleanly separating the few likely pathogenic variants from the much larger background of neutral missense variants.

## Results

### Searching for regional missense constraint within transcripts

We used a set of 17,915 transcripts (see Materials and Methods for transcript filtering; **Table S2**) and, for every exon, extracted rare (minor allele frequency [MAF] < 0.1%) missense variants from the Exome Aggregation Consortium (ExAC; n = 60,706)^13^ dataset and predicted the expected number as described previously^15^. In our earlier work, we identified 1614 transcripts that were significantly (Z ≥ 3.09;p < 10^-3^) depleted of missense variation^13^. To move away from statistical significance and towards biological significance, here we focus on the fraction of expected missense variation that is observed (defined as γ).

Given that missense deleteriousness depends partially on the location of the variant within the transcript, we applied a likelihood ratio test to determine if γ was uniform along a transcript or whether that transcript had evidence of distinct domains of missense constraint. The method is depicted in **Figure S1** and explained in detail in the Materials and Methods. We found evidence of significant regional differences in missense depletion within 2700 transcripts (15.1%) with 1717 split into two segments (having one significant break), 904 with three regions, and 79 with four or more distinct regions of missense tolerance (**Table S3** for distribution data; **Table S4** for regional constraint data). The transcripts with regional missense constraint have, on average, more coding base pairs (bp) than all other transcripts (2753 versus 1530 bp; Wilcox p < 10^-50^), which is expected given the increased power to detect regional patterns in missense variation in longer transcripts.

For all following analyses, we combined whole transcripts and partial segments of transcripts with the same values of γ when exploring disease relevance. As an example, ~14% of the coding region has γ < 0.6; it is compromised of 1789 unbroken transcripts and 2996 segments from transcripts with distinct regions of missense constraint.

### ClinVar variants and regional depletion

We previously found that missense depleted genes were enriched for known disease genes in the Online Mendelian Inheritance in Man (OMIM) database^15^, and therefore hypothesized that the missense depleted regions of genes would be similarly enriched for disease-associated variation. To test this hypothesis, we extracted pathogenic variants from ClinVar^25^ to evaluate any potential enrichments. Since our method focused on finding regions that are intolerant of heterozygous missense variants, we selected only those variants that disrupted haploinsufficient genes known to cause severe disease (n = 49 genes with 404 variants; **Tables S5-6**). Transcripts and regions of transcripts with the fraction of expected variation observed (γ) ≤ 0.6 contain 89% of all the ClinVar pathogenic missense variants (6.2-fold enriched; p < 10^-50^; **Table S7**), despite only representing 14% of the coding region of the human genome.

Of the 49 severe haploinsufficient genes, 32 (65%) have evidence of regional variability in missense constraint, and of this subset, 22 (69%) contain both constrained and unconstrained regions. As an example, the first 283 amino acids of *CDKL5* have only 24% of the expected number of missense variants (χ^2^ = 37.6), while the remaining 748 amino acids in the gene have 82% (χ^2^ = 5.5). ClinVar lists 43 pathogenic or likely pathogenic missense variants in *CDKL5*, 39 (91%) of which reside in the constrained regions (**Figure 1**). Three of the remaining variants are in the first 50 base pairs (bp) of exon 10, just outside the boundary defined by our approach, and lie in the kinase domain that extends 66 bp into that exon.

**Figure 1.**
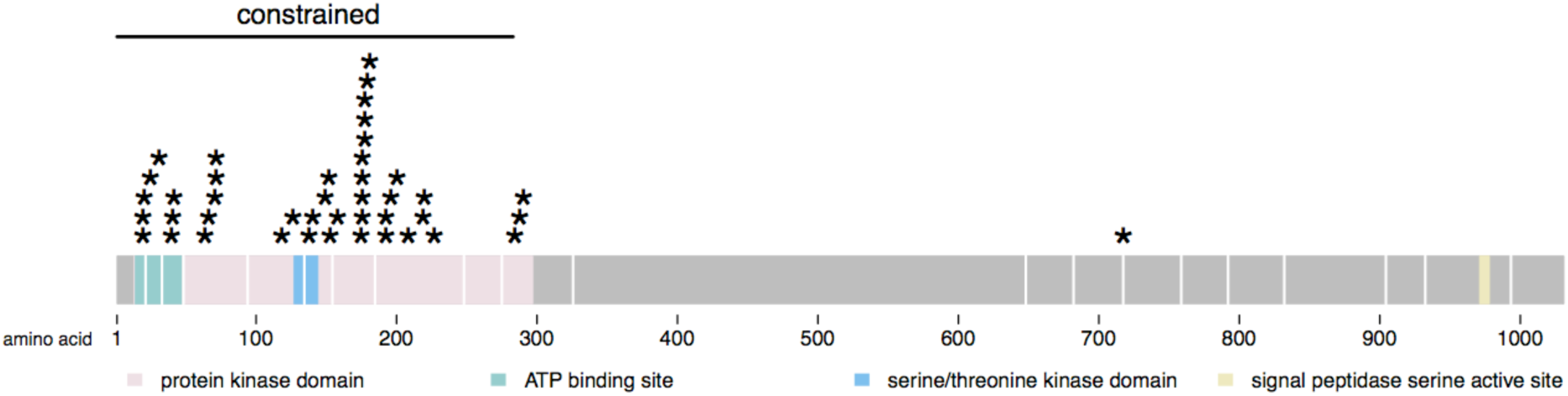
Distribution of ClinVar^25^ pathogenic and likely pathogenic variants in *CDKL5*. Pathogenic variants from ClinVar are indicated with a star. 91% of the variants (39/43) fall into the first 283 amino acids, which are significantly constrained (24% of expectation observed, χ^2^ = 37.6). The constrained region is marked with a bar.

### Using regional constraint to interpret *de novo* variation

The variants we tested from ClinVar are considered pathogenic, but we wanted to evaluate the ability of our regional missense depletion results to aid in prioritization of variants where an unknown subset of the variants are likely contributing to disease. For this purpose, we chose to study *de novo* missense variants from cases with a neurodevelopmental disorder (n = 5620; **Table S8**)^1^^-^^6^ due to the significant, but modest, excess of *de novo* missense variants in these cases (1.3-fold enriched; p < 10^-50^; **Table S1b**). The *de novo* missense variants from 2078 unaffected siblings of autism spectrum disorder cases were used as controls (**Table S8**)^10,11^.

As depicted in **Figure 2a**, the distribution of control *de novo* missense variants between bins of missense depletion closely matches the distribution seen for coding base pairs overall. To illustrate, 87.9% of the control variants reside in regions with γ > 0.6, which represent 85.6% of all coding bases (Chi-squared test p = 0.1262). By contrast, the *de novo* missense variants identified in patients with a neurodevelopmental disorder are enriched in the most missense-depleted regions (2.0-fold enriched in regions with γ ≤ 0.6, Chi-squared test p < 10^-50^).

**Figure 2.**
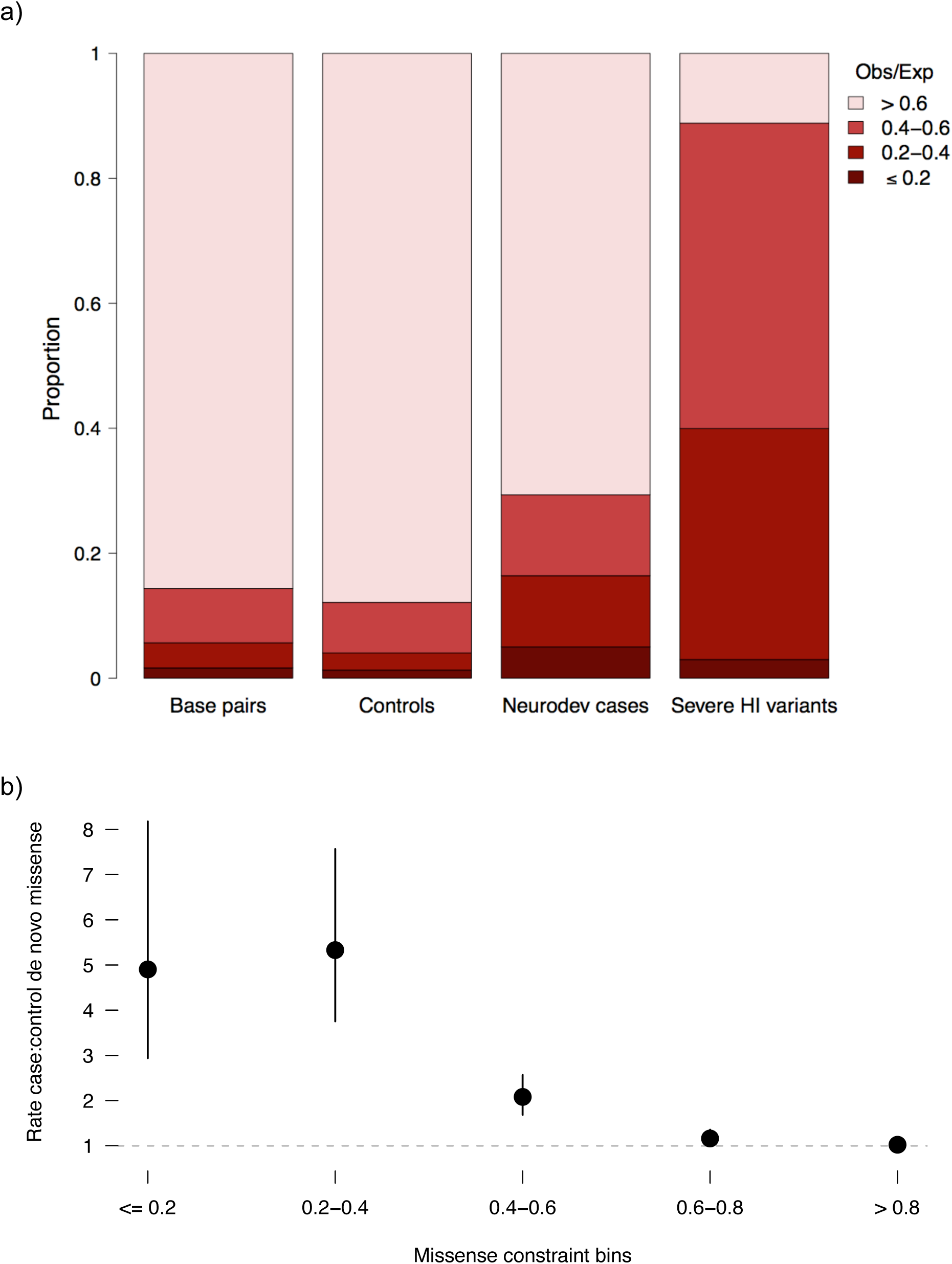
Missense depleted regions of the exome are enriched for established pathogenic variants and *de novo* missense variants found in patients with neurodevelopmental disorders. **a**) Depicted for each bin of missense depletion (e.g. fraction of expected variants observed [γ] > 0.6) is the fraction of coding base pairs (Base pairs), *de novo* missense variants from 2078 control trios (Controls)^10,11^, *de novo* missense variants from 5620 cases with a neurodevelopmental disorder (Neurodev cases)^1^^-^^6^, and pathogenic or likely pathogenic missense variants from ClinVar^25^ in haploinsufficient genes that cause severe disease (Severe HI variants). Darker reds indicate more extreme missense depletion. **b**) Comparison of the rate of case *de novo* missense variants to the corrected rate of control *de novo* missense variants by bins of missense depletion. Here, we have added a bin for 0.6 < γ ≤ 0.8. The case variants come from 5620 trios with a neurodevelopmental disorder^1^^-^^6^ and the control variants were identified in 2078 control trios^10,11^. The control rate was corrected by the ratio of the rate of *de novo* synonymous variants in cases versus controls (~1.14). The dashed gray line indicates a ratio of one. 95% confidence intervals are depicted around each point estimate.

We next wanted to compare the rates of *de novo* missense variants between the neurodevelopmental disorder cases and the controls. Since these datasets were sequenced at separate times and locations, we first needed to correct for any potential differences that might result in different variant ascertainment rates between datasets. In particular, we note that the cases were more recently sequenced than the controls, and consequently exhibit a significantly higher rate of synonymous variation (0.287 *de novo* synonymous variants per case exome versus 0.251 per control exome, two-sided Poisson exact test p = 9.80x10^-3^). We control for this difference in synonymous rate by multiplying all rates of *de novo* variation in the controls by the ratio of the synonymous rate in the cases compared to controls (~1.14). This correction will reduce apparent signal, but ensures that our results are not inflated by technical differences in mutation discovery.

We compared the rate of *de novo* missense variants in cases to the corrected rate in controls across four bins of missense constraint. If a region or transcript is tolerant of missense variation, we expect it to have the same rate of *de novo* variation in cases as in controls, reflecting the background rate of mutation (ratio of 1:1). However, if the region is intolerant of missense variation—and therefore more likely to be associated to disease—we expect to observe a higher rate of *de novo* variants found in cases compared to controls (ratio >1:1). Here, we have added a bin for those genes and regions with 0.6 < γ ≤ 0.8. Both of the least missense-constrained bins (γ > 0.6) are indistinguishable from 1, supporting our choice of 0.6 as a cut-off between constrained and unconstrained regions (**Figure 2b; Table S9**). Combined, there are 0.58 *de novo* missense variants per case exome and 0.56 per control exome after correction for the difference in the rate of *de novo* synonymous variation.

By contrast, the three most depleted bins together (γ ≤ 0.6) have 0.24 *de novo* missense variants per case exome and only 0.08 per control exome after correction. There is quantitative value in the missense depletion parameter, however, as suggested by the intermediate OR of regions and genes with more modest missense depletion (0.4 < γ ≤ 0.6; OR = 2.1) compared to the most depleted two bins (γ ≤ 0.4; OR > 4.9; **Figure 2b; Table S9**). It is important to note that the majority (71%) of *de novo* missense variants found in cases reside in transcripts and regions that are not missense constrained (γ > 0.6). However, we have used these analyses to refine the signal of *de novo* missense variant enrichment and, by doing so, shrunk the number of candidate pathogenic variants from 4683 to 1374.

Taken together, and consistent with related observations regarding truncating variation, these analyses indicate that the signal for both established pathogenic variants as well as the excess of *de novo* missense variants in cases with a neurodevelopmental disorder can be found in those transcripts and regions with 60% or less of their expected missense variation.

### Measuring the increased deleteriousness of amino acid substitutions

While the general mutational tolerance of a gene or region disrupted by a missense variant carries significant information, it is also critical to consider the specific type of amino acid substitution that occurred. Major changes in the physiochemical properties of the amino acid side chain are expected to have larger effects on the protein than more subtle changes to the side chain. The deleteriousness of these changes has been quantified in a variety of metrics, the two most common of which are BLOSUM^24^ and Grantham^23^. Here, we hypothesized that there might be specific amino acid substitutions that are preferentially eliminated when they occur in the most missense depleted regions of the human exome. That is, analogous to the way we recognize the relative tolerance of genes to mutation by their variation content, we can potentially compare the disruptiveness of different amino-acid substitutions by how tolerated they are across all genes.

To measure the increased deleteriousness of amino acid substitutions when they occur in the constrained regions of the exome, we tabulated all possible amino acid-to-amino acid substitutions that could occur in the exome via a single nucleotide mutation along with the number observed in ExAC (with MAF < 0.1%)^13^. The rate of possible substitutions observed was determined for constrained (γ ≤ 0.6) and unconstrained (here, γ > 0.8) regions separately; in almost all instances, we observed a higher rate in the unconstrained regions, including for synonymous variants. The fold difference between the rate in the unconstrained and constrained regions differs depending on the functional class. For synonymous changes, the rate difference clusters around one and enters the 3-5 range for nonsense changes. As expected, the rate difference for missense changes falls primarily in between the synonymous and nonsense changes (**Figure S2**).

We used the normalized fold difference of missense substitutions (“missense badness”) as a measure of the increased deleteriousness of amino acid substitutions when they occur in constrained genes and regions (scores provided in **Table S10**). As expected, this score has a high correlation with both BLOSUM and Grantham scores (r = −0.6437 and 0.5180, respectively; **Figure S3**) with few exceptions.

### Combining variant level deleteriousness scores

We wanted to determine which variant deleteriousness metric, or combination of metrics, best differentiated benign from pathogenic missense variants. We selected missense variants with a minor allele frequency (MAF) > 1% in ExAC as our benign set (n = 82,932 variants after removing those variants missing one of the metrics) and used the ClinVar missense variants found in haploinsufficient genes that cause severe disease as our set of pathogenic variants (n = 402 after removing variants missing one of the metrics). We compared the following metrics: missense depletion (γ) of the region in which the variant resides, missense badness, PolyPhen-2^19^, BLOSUM^24^, and Grantham scores^23^. Using a series of logistic regressions, missense depletion (γ) of the region in which the variant was located best predicted missense deleteriousness (**Table S11**).

As the metrics can provide complementary information, we sought to create a composite predictor. Given that γ was by far the best score, we tested nested logistic regression models and found that both missense badness and PolyPhen-2 significantly added to the composite predictor, but neither BLOSUM nor Grantham did. Therefore, the best model included γ, missense badness, and PolyPhen-2 (**Table S12**), and we took the predictions as our final score, known as MPC (for **M**issense badness, **PolyPhen-2**, and **C**onstraint). Given the number of benign variants, the range of MPC is limited from 0 to 5, with larger numbers indicating increased deleteriousness.

### Using MPC to evaluate the deleteriousness of *de novo* variants

To independently test the utility of these scores, we analyzed the MPC distributions of the previously described *de novo* variants from 5620 cases with a neurodevelopmental disorder^1^^-^^6^ and from 2078 controls^10,11^. We find that the vast majority (72%) of *de novo* missense variants in controls have MPC values below 1 (**Figure 3**). In fact, those *de novo* missense variants with MPC < 1 show no sign of enrichment in cases compared to the rate seen in controls (0.507 *de novo* missense variants per case exome versus 0.504 in controls after correction; rate ratio = 1.01; one-sided Poisson exact p = 0.885; **Table 1**). By contrast, the MPC distribution for the *de novo* missense variants identified in cases with a neurodevelopmental disorder appear to comprise two distributions: one following the distribution of the control *de novo* variants and the other with a peak at an MPC of 2.5 (**Figure 3**), reinforcing that these variants are a mix of signal and noise. Variants with MPC ≥ 2 have a rate nearly 6 times higher in cases than in controls (0.162 versus 0.028 *de novo* missense variants; rate ratio =5.79; one-sided Poisson exact p < 10^-50^; **Table 1**), while those with intermediate MPC values (1 ≤ MPC < 2) have a more modest excess in cases (0.185 per case versus 0.124 per control exome after correction; rate ratio = 1.49; one-sided Poisson exact p = 3.70x10^-9^; **Table 1**).

**Figure 3.**
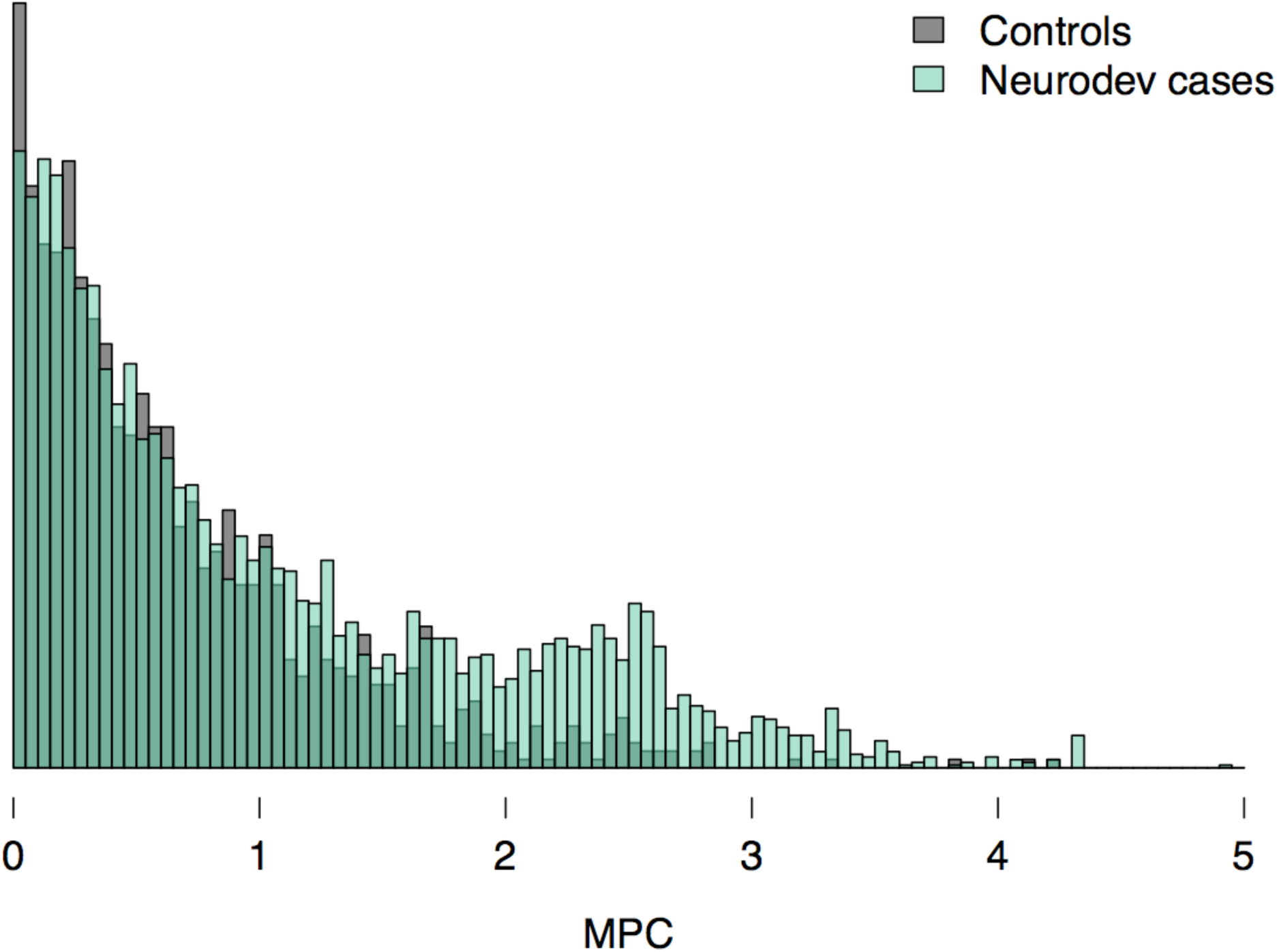
The MPC distributions for *de novo* variants in cases with a neurodevelopmental disorder and controls. The MPC scores for *de novo* missense variants identified in 2078 controls^10,11^ are depicted in gray. Note that the control rate has been corrected for the differences in *de novo* synonymous rates between cases and controls. In green are the MPC scores for the *de novo* missense variants (n = 5113) identified in 5620 patients with a neurodevelopmental disorder^1^^-^^6^. Larger MPC values indicate increasing deleteriousness of the variant.

**Table 1.** Distribution of MPC scores for *de novo* missense variants in patients with neurodevelopmental disorders and controls. For three bins of MPC values, the number of *de novo* missense variants ( count) and per exome rate are reported for 5620 cases with a neurodevelopmental disorder^1^^-^^6^ and 2078 controls^10,11^. Below each bin is the rate ratio. Higher MPC values indicate increasing deleteriousness. * The per exome rate for the controls has been corrected for the difference in rate of *de novo* synonymous variants in cases versus controls (~1.14:1). P-values come from one-sided Poisson exact tests.

|  |  | **MPC < 1** | **1 ≤ MPC < 2** | **MPC ≥ 2** |
| --- | --- | --- | --- | --- |
| **Neurodevelopmental disorder cases** | Count | 2848 | 1038 | 911 |
|  | Per exome | 0.507 | 0.185 | 0.162 |
| **Controls** | Count | 918 | 226 | 51 |
|  | Per exome* | 0.504 | 0.124 | 0.028 |
|  | **Rate ratio**<br>**p-value** | 1.01<br>0.885 | 1.49<br>3.70x10<sup>-9</sup> | 5.79<br><10<sup>-50</sup> |

We compared MPC to three other metrics that evaluate the deleteriousness of missense substitutions: PolyPhen-2^19^, CADD^20^, and M-CAP^22^. We combined the scores for *de novo* missense variants from cases and controls and took the top ~10% ranked by each score to determine the enrichment of case variants. Since the total proportion of case variants is 0.8 (5113 out of 6382), if a metric has no predictive value, the proportion of case variants in the top 10% should match the overall rate (0.8). However, the better a metric is at determining deleteriousness of variants, the greater the fraction of case variants should be seen in the top 10% of scores – since the global excess of mutations in cases suggests up to 20% of *de novo* missense variants may confer risk to neurodevelopmental disorders. While all four metrics are significantly enriched for case *de novo* variants in their most deleterious ~10% of variants, MPC has the greatest enrichment of cases variants (odds ratio [OR] = 5.43; Fisher’s exact test p < 10^-27^; **Table 2**; **Table S13**). Given that PolyPhen-2 is incorporated into each of MPC, CADD, and M-CAP, it is unsurprising that PolyPhen-2 by itself shows the least enrichment (OR = 1.44; Fisher’s exact test p = 2.66x10^-3^; **Table 2**; **Table S13**), though clearly it is an integral component of the predictive success of the other metrics. The greater enrichment for MPC supports the notion that high MPC values are more specific to pathogenic variants than high CADD, M-CAP, or PolyPhen-2 scores. We provide MPC values for all possible missense variants in the canonical transcripts under study in this work (**Table S14**).

**Table 2.** Comparison of MPC, M-CAP, CADD, and PolyPhen-2 scores for *de novo* missense variants in patients with neurodevelopmental disorders and controls. Taking the combined scores from 5620 cases with a neurodevelopmental disorder^1^^-^^6^ and 2078 controls^10,11^, we took the top ~10% of scores and determined the fraction of those variants that came from the cases. We also report odds ratios and p-values (Fisher’s exact test) for the enrichment of case variants in the top ~10%. Overall, the proportion of case variants is 0.80. See **Table S13** for more details.

|  | **MPC** | **M-CAP** | **CADD** | **PolyPhen-2** |
| --- | --- | --- | --- | --- |
| Fraction of top 10% from cases | 0.95 | 0.93 | 0.86 | 0.85 |
| Odds ratio | 5.43 | 3.52 | 1.58 | 1.44 |
| p-value | $1.48 \times 10^{-28}$ | $4.35 \times 10^{-20}$ | $1.46 \times 10^{-4}$ | $2.66 \times 10^{-3}$ |

## Discussion

We have developed a method to locate regions within genes that are specifically intolerant of missense variation and used those results to create MPC, a metric to evaluate the deleteriousness of missense variants. Our work here highlights the usefulness of incorporating regional depletion of missense variation into missense variant pathogenicity calculations.

While many genes have a relatively consistent tolerance or intolerance to missense mutation throughout, ~15% display evidence of strong regional variability in missense constraint and are split into two or more regions of varying missense depletion. We find that the genes and regions that have 60% or less of their expected missense variation—while only representing a small fraction of all coding sequence— contain 89% of missense pathogenic variants^25^ in haploinsufficient genes known to cause severe disease. These genes and regions also contain the vast majority of the excess of *de novo* missense variation that is seen in 5620 cases with a neurodevelopmental disorder^1^^-^^6^.

Since the missense constrained regions are depleted of variation due to selective pressures, we proposed that including information about the local missense depletion could improve variant deleteriousness metrics. We first created a measure of the increased deleteriousness of amino acid substitutions when they occur in missense constrained genes and regions, which outperformed similar amino acid substitution matrices (BLOSUM^24^ and Grantham^23^) at separating pathogenic^25^ from benign variants. The best predictor of variant deleteriousness, however, was the combination of regional missense constraint, the amino acid substitution score we developed (missense badness), and PolyPhen-2^19^. The MPC scores—the joint metric—for the *de novo* missense variants from neurodevelopmental cases appeared to comprise a mixture of two distributions (benign and pathogenic), which matches what would be expected given the modest enrichment of such variants in the cases.

When applied to the *de novo* missense variants identified in 5620 patients with a neurodevelopmental disorder and 2078 controls, MPC displayed the greatest specificity for pathogenicity when compared to three other metrics of variant deleteriousness. Specifically, we found that *de novo* missense variants with MPC ≥ 2 were found at a 5.8 times higher rate in cases compared to controls. This enrichment approaches that found for *de novo* protein-truncating variants—which are predicted to be more deleterious overall than missense—that were absent in the ExAC database and that disrupted genes extremely intolerant of protein-truncating variation (pLI ≥ 0.9; rate ratio = 6.7)^17^. We predict that MPC will be most informative for those variants that are found in regions with intermediate missense depletion (40-60% of expected variation), since this set of variants has a lower signal to noise ratio than the variants found in the more missense depleted genes and regions.

Ideally, constraint would be calculated per base, but even the 60,706 individuals released in the original ExAC dataset is not large enough to provide sufficient power to do this. We therefore needed to aggregate variant counts and, while there are many options, we chose to aggregate across exons with further refinement to the amino acid level. As publicly released datasets such as ExAC continue to grow, our ability to define the constraints on specific amino acids and base pairs will improve.

Moving forward, it will also be critical to consider both noncoding DNA and nonlinear coding sequences. Currently, determining noncoding regions under selective constraint is limited by our lack of knowledge of both boundaries of relevant elements and which variants are more likely to be deleterious, both of which would decrease power to identify intolerant elements^26^. However, we can start to evaluate non-linear sequences, which would be a critical advance. Binding pockets, which are key functional domains of proteins, are made up of amino acids scattered across the gene and are currently not evaluated by our method. Other 3D structural aspects of the protein (e.g. internal versus external residues, structured versus unstructured regions) could also supply important information to consider when evaluating variant deleteriousness. Evaluating disparate amino acids would also provide the ability to determine constraint of the same residue or set of residue across members of a protein family. Therefore, future work would greatly benefit from being able to analyze nonlinear sequences. The knowledge gained from our work and similar studies will continue to improve our ability to interpret genetic variation and, therefore, understanding of the genetic basis of disease.

## URLs

Exome Aggregation Consortium (ExAC), http://exac.broadinstitute.org/; ClinVar https://www.ncbi.nlm.nih.gov/clinvar/; Online Mendelian Inheritance in Man (OMIM), http://omim.org/; ClinGen Dosage Sensitivity Map www.ncbi.nlm.nih.gov/projects/dbvar/clingen/; Combined Annotation Dependent Depletion (CADD) http://cadd.gs.washington.edu

## Acknowledgements

We would like to thank all members of the Analytic and Translational Genetics Unit for feedback and guidance. In particular, we thank Alex Bloemendal for his help with mathematical annotation and Eric Minikel for providing the ClinVar variant list as part of Lek et al 2016. Funding for this work came from 5U01MH094432-02, 1R01MH109539- 01, and SCSBMIT. KJK is supported by an NIGMS Fellowship (F32GM115208). AHODL is supported by a NIGMS training grant (T32GM007748).

## Author Contributions

KES, BMN, and MJD conceived and designed the experiments. KES executed the analyses. JAK, KJK, AHODL, and EPH provided data and analysis suggestions. DGM, BMN, and MJD supervised the research. KES and MJD performed the primary writing. All authors approved of the final manuscript.

## Competing Financial Interests

The authors declare no competing financial interests.

## Online Material and Methods

### Transcript and exon definitions

In order to have one representative transcript for each gene, we used the canonical GENCODE (v19) transcript as defined by Ensembl 75, for protein-coding genes. We removed transcripts that lacked a methionine at the start of the coding sequence, a stop codon at the end of coding sequence, or were indivisible by three, which left 19,621 transcripts. Additionally, we dropped 795 transcripts that had zero observed variants when dropping counts in exons with a median depth < 1. Our previous work with the Exome Aggregation Consortium’s dataset (ExAC; n = 60,706)^13^ identified a set of 251 genes that had either (1) far too many synonymous and missense variants as determined by a Z score (p < 10^-4^ and 10^-3^, respectively) or (2) far too few synonymous and missense variants as determined by a Z score (p < 10^-4^ and 10^-3^, respectively). These outlier genes were removed as well as all genes with synonymous Z scores that were significantly high or significantly low (p < 10^-3^; n = 310), leaving 17,915 transcripts for all analyses (**Table S2**). The exon boundaries were defined by UCSC’s annotation for GENCODE v19 (downloaded on June 16^th^, 2014).

### Observed variant counts

To obtain the observed number of missense variants per exon, we extracted variants from ExAC that met the following criteria:

1. Defined as a missense change by the predicted amino acid substitution. Variants that would be considered “initiator_codon_variants” and “stop_lost” by annotation programs such as Variant Effect Predictor (VEP)^27^ are therefore included in the total.
2. Caused by a single nucleotide change.
3. Had an adjusted allele count ≤ 123, corresponding to a minor allele frequency (MAF) < 0.1% in ExAC. The adjusted allele count only includes individuals with a depth (DP) ≥ 10 and a genotype quality (GQ) ≥ 20.
4. Had a VQSLOD ≥ −2.632.

Due to the VQSLOD threshold, variants were not required to have a PASS in their FILTER column. The observed counts represent the unique number of qualifying variants and not the aggregate allele count of all qualifying variants within the exon. Variants in exons with median depth < 1 were not considered in our analyses.

### Expected variant counts

Expected missense variant counts were determined as described in Lek et al.^13^ Briefly, we used a model of mutation based on sequence context and corrected for regional divergence between humans and macaques to define the probability of a mutation per exon in all canonical transcripts^13,15^. We used exons with a median depth ≥ 50 and regressed the number of rare, synonymous variants on the probability of a synonymous mutation. Regressions were run separately for the autosomes with the pseudo-autosomal regions (PAR) of the X chromosome, the non-PAR regions of the X chromosome, and the Y chromosome. The expectations produced by these regressions were then corrected for the median depth of coverage of the exon using the following equation:

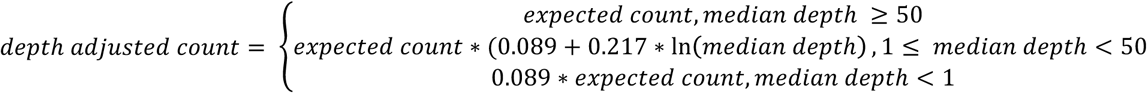

For exons with a median depth < 1, we set the expected counts to 0.

### Likelihood ratio tests to define regional constraint

Using the observed and expected counts for the 17,915 canonical transcripts, we searched for significant breaks between exons that would split the transcript into two or more regions with varying levels of missense depletion. For these analyses, we assume that observed counts should follow a Poisson distribution around the expected number. We defined the null model—no regional variability in missense depletion—as the model where the overall fraction of expected missense variation observed (*γ*) for the transcript is used as the expectation for all segments. We then employed a likelihood ratio test to compare the null model with an alternative model where expectation was *γ* for each specific segment. Given that the alternative model should always have a better fit than the null, we require a χ^2^ above a given threshold to establish significance.

We used the following general formula to determine the significance of a break that would split a transcript into segments *A* and *B*:

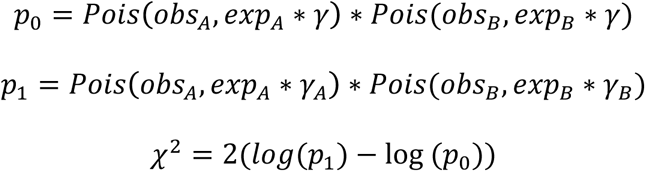

Where *γ* is the fraction of expected variation observed across all segments in the transcript; *obs_A_* is the observed number of missense variants in segment *A*; *exp_A_* is the expected number of variants in segment *A*; *γ_A_* is the fraction of expected variation observed only for segment *A*; *obs_B_* is the observed number of missense variants in segment *B*; *exP_B_* is the expected number of variants in segment *B*; *γ_B_* is the fraction of expected variation observed only for segment *B*; and *Pois* denotes the Poisson likelihood.

To account for the fact that exon boundaries do not always follow the boundaries of biological significance, we refined any significant break we found by searching for an amino acid nearby that best modeled the differences in missense depletion. Specifically, we tested all amino acid boundaries halfway through each of the bordering exons. As an example, if the 5’ exon for a significant break was 100 amino acids long, we tested each amino acid break for the 50 amino acids at the 3’ end of the exon (and therefore closest to the original exon break). This would be repeated for half of the amino acids in the 3’ exon as well. The amino acid boundary that best modeled the data (via the χ^2^) was taken as the break between the two regions of the gene.

For the purposes of this method, all exons or sections with more observed variants than expected were assigned *γ* = 1 since we were looking for variation in missense depletion. In addition, exons or sections with zero observed variants were considered to have one variant to prevent *γ* = 0.

We first searched for a single break in between exons that would significantly (χ^2^ ≥ 10.8, p < ~10^-3^) better model the transcript’s data than the null model. If multiple significant breaks between exons were found, we took the best break as defined by the χ^2^ value. If a significant break was found, we first refined the break by examining the local amino acid space and then searched for a second break. This process was repeated until the best break did not significantly improve on the model (χ^2^ < 10.8). If a transcript had no significant single break, we searched for two breaks at a time, requiring a χ2 ≥ 13.8 (p < ~10^-4^) to indicate significance. As before, if we found evidence of two breaks, both breaks were refined to the local amino acids that gave the most significant χ^2^. Those transcripts with χ2 < 13.8 were considered to show no evidence of regional variability in missense depletion, and were left intact. The general process is depicted in **Figure S1** and the regional constraint data can be found in **Table S4**.

### ClinVar pathogenic variants

We extracted variants from the July 9, 2015 release of ClinVar^25^ that were labeled as “pathogenic” and “likely pathogenic”. We specifically focus on those missense variants that fell into a set of 55 haploinsufficient genes (49 when removing outliers; **Table S5**) that cause severe disease (n = 483 variants; 404 variants when removing synonymous Z-score outliers; **Table S6**). The haploinsufficient genes were primarily those with sufficient evidence for dosage pathogenicity (level 3) as determined by the ClinGen Dosage Sensitivity Map <u>(www.ncbi.nlm.nih.gov/projects/dbvar/clingen/;</u> downloaded on May 5, 2015); the severity of disease caused by variants in the genes was manually curated by AHODL. We also included 11 genes that were established to cause severe disorders in a dominant and/or haploinsufficient mechanism, as manually curated by AHODL and EPH.

### *De novo* variants from cases with a neurodevelopmental disorder

We collected the *de novo* variants found in 5264 trios with intellectual disability/developmental delay^1^^-^^5^ and 356 with an epileptic encephalopathy^6^. *De novo* variants from the unaffected siblings of autism cases were used as controls (n = 2078)^10,11^. As described in Kosmicki et al.^17^, we created a standardized annotation of all *de novo* variants to ensure uniformity across the datasets. Briefly, we used a Python implementation of vt normalize^28^ to annotate all variants with VEP^27^ version 81 with GENCODE v19 on GRCh37. Given that some variants fall into multiple transcripts, we used the annotation from the canonical transcript whenever possible. In the case of multiple canonical transcripts or no canonical transcript available for the gene, the most deleterious annotation was used. All variants are provided in **Table S8**.

### Correcting the rate of de novo variants in controls

Before we could compare the rate of *de novo* missense variants between the cases and controls, we wanted to correct for potential differences in sequencing technology and time of sequencing (e.g. the cases were sequenced more recently than the controls). As listed in **Table S1b**, the rate of *de novo* synonymous variants in the cases (0.287 events per case exome) is significantly higher than the rate seen in controls (0.252 events per control exome; p = 9.80x10^-3^). This increased in rate would also extend to missense variation, and therefore make any enrichment using the raw rates of *de novo* missense variants inflated. To correct for this, we multiplied all control *de novo* rates in analyses by ~1.14, which is the ratio of the rate of *de novo* synonymous variants in cases to the rate seen in controls. Our correction for this difference is conservative, as it will reduce any signal of enrichment we see in cases.

### Confidence intervals around the ratio of case:control *de novo* variant rates

We compared the rate of *de novo* missense variants in cases compared to the rate in controls for the five constraint bins. To determine confidence intervals around the point estimates of the ratio of *de novo* variant rates, we took the natural logarithm of the point estimate

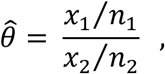

and found the standard error

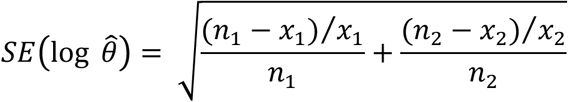

using the delta method. The upper and lower bounds are then transformed back to obtain the 95% confidence interval

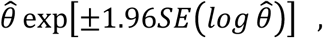

where *x*_1_ is the number of case *de novo* variants; *n*_1_ is the number of case trios; *x*_2_ is the number of control *de novo* variants; and *n*_2_is the number of control trios.

### Creation of missense badness

We created a metric (missense badness) of the increased deleteriousness of specific amino acid substitutions when they occur in constrained regions to identify those substitutions that are preferentially eliminated when they occur in missense depleted sequence. To do so, we identified all possible amino acid-to-amino acid substitutions that could occur via a single nucleotide mutation and then tallied the number of these substitutions in ExAC with a MAF < 0.1%. The observed and possible were then split by whether they occurred in a gene or regions with γ ≤ 0.6 (constrained) or γ > 0.8 (unconstrained) and we determined the rate of possible substitutions observed for both groups. Transcripts and regions with 0.6 < γ ≤ 0.8 were excluded from both constrained and unconstrained groups because, while these regions look null with respect to the rate of *de novo* missense variants in cases compared to the rate seen in controls (**Figure 2b**), they are noticeably depleted of the expected amount of missense variation and may hold some small residual signal. We observed a higher rate of possible substitutions observed in the unconstrained regions with the notable exceptions that synonymous changes in isoleucine and phenylalanine did not follow this pattern.

We used the median fold difference of all synonymous substitutions as a floor (set to 0) and the median of all nonsense substitutions as a ceiling (set to 1) and normalized the missense fold differences to create missense badness. Missense badness is provided in **Table S10**.

### Creation of MPC, a composite missense deleteriousness score

We used logistic regressions to determine which of five deleteriousness metrics was best at separating benign from pathogenic missense variants. The metrics we compared were the missense depletion of the region in which the variant was found (γ), missense badness, PolyPhen-2^19^, BLOSUM^24^, and Grantham scores^23^. Our benign variants were missense variants with a MAF > 1% in ExAC^13^ (n = 88,083 variants when removing synonymous Z outlier genes). The pathogenic variants were ClinVar^25^ missense variants found in haploinsufficient genes that cause severe disease (n = 404 variants when removing synonymous Z outlier genes).

As the metrics provide complementary information, we used nested models to determine the best composite score starting with missense depletion (γ). Missense badness and PolyPhen-2 significantly added to the composite predictor, but BLOSUM and Grantham did not. We therefore tested the combination of the three significant metrics and all possible interactions between them. The best model included all three scores and the interaction between γ and missense badness as well as the interaction between γ and PolyPhen-2 (**Table S11**).

We used the best regression to predict scores for all benign and pathogenic variants. In order to make more easily interpretable numbers, we transformed the raw score (*RS*)

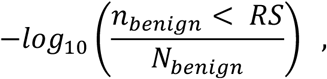

Where *n_benign_* is the number of benign variants with a raw score less than *RS* and *N_benign_* is the total number of benign variants. We refer to the final composite score as MPC. Since there are ~88k benign variants that had information for all three metrics, the highest MPC is ~5. MPC values for all possible missense variants—excluding those involving a selenocysteine—in the 17,915 canonical transcripts were calculated and are provided in **Table S14**.

